# Bifurcation analysis of an impulsive system describing Partial Nitritation and Anammox in a hybrid reactor

**DOI:** 10.1101/2020.02.14.949099

**Authors:** Matthew J. Wade, Gail S. K. Wolkowicz

## Abstract

Low-energy deammonification under mainstream conditions is a technology that has received significant attention in recent years as the water industry drives towards long-term sustainability goals. Simultaneous partial nitritation-Anammox (PN/A) is one process that can provide substantial energy reduction and lower sludge yields. Mathematical modelling of such a process offers engineers insights into the conditions for maximising the potential of PN/A. Laureni *et al., Water Res*. (2019) have recently published a reduced mechanistic model of the process in a sequencing batch reactor, which indicates the effect of three key operating parameters (Anammox biofilm activity, dissolved oxygen concentration and fraction of solids wasted) on performance. The analysis of the model is limited, however, to simulation with relatively few discrete parameter sets. Here, we demonstrate through the use of bifurcation theory applied to an impulsive system, that a phase space can be generated describing the continuous separation of system equilibria. Mapping process performance data onto these spaces allows engineers to target suitable operating regimes for specific objectives. Here, for example, we note that the nitrogen removal efficiency is maximised close to the trans-critical bifurcation curve denoting nitrite oxidising bacteria washout, but control of solids washout and Anammox biofilm activity can also reduce oxygen requirements whilst maintaining an appropriate Hydraulic Retention Time. The approach taken is significant given the possibility for using such a methodology for models of increasing complexity, which will enable engineers to probe the entire parameter space of systems of higher dimensionality and realism in a consistent manner.

## 1 Introduction

The deammonification of wastewater is a well-studied process that nevertheless remains an area for intensive research given the continued drive towards energy and resource sufficiency. Technologies and processes that offer equivalent or better nitrogen removal efficiencies at lower cost or energy consumption have become a principal research focus for stakeholders in the wastewater industry. Partial nitritation coupled to anaerobic ammonium 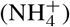 oxidation (Anammox), PN/A, presents a means to shortcut the conventional nitrification-denitrification treatment pathway, reduce oxygen and carbon requirements, and minimise sludge production [14, 4].

Mathematical modelling of the PN/A process has the potential to offer engineers qualitative insights into operating regimes relating to reactor configuration and control. Modelling of such systems has generally been restricted to deterministic Ordinary Differential Equation (ODE) models, although one-dimensional Partial Differential Equations (PDEs) that consider nutrient gradients through the biofilm have been investigated in the context of control [28] and as component of larger biological wastewater treatment models as part of commercial software [27]. An extension of the widely-used Activated Sludge Models (ASMs) to account for nitrite dynamics with the inclusion of biomass terms for the Ammonium and Nitrite Oxidising Organisms (AOB and NOB) as separate state variables [12] has motivated their use for simulation and control of PN/A systems [23, 29, 30, 28, 10]. Others have applied modelling to address specific process performance questions in floc-based [19], granular sludge [1] or biofilm systems [7, 18], or to explore fundamental questions related to microbial ecology, such as coexistence in nitrifying systems [2] and competition [20, 22].

Recently, attempts have been made to model the PN/A process in hybrid systems (mixed floc/granule and attached biofilms) [9, 16], indicating that process performance is highly sensitive to process design (e.g. degree of flocculation and segregation) and operation (e.g. organic loading rate). These findings, coupled with earlier empirical work on understanding stability in PN/A system [11, 15] suggest that exploring the process with adequate models and rigorous mathematical analysis could result in a better understanding of its behaviour and stability under a range of process configurations.

The slow growth kinetics of Anammox bacteria (AMX) led to the early adoption of Sequencing Batch Reactors (SBR) as a means to retain this critical biomass, whilst maintaining homogeneous and stable operation of the system [21]. The relatively simple model proposed by Laureni *et al*. [16] describes PN/A in a hybrid moving bed biofilm reactor (MBBR) as a mainstream, rather than conventional sidestream, process. At lower mainstream temperatures, around 15°C, the kinetic advantage that Ammonium Oxidising Organisms AOB had over the NOB is known to be diminished, with comparable growth rates observed experimentally [8]. The objective of the model was to understand the operating conditions required to suppress NOB activity and achieve complete comproportionation of ammonium and nitrite 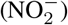, to dinitrogen gas (N_2_), through the Anammox shortcut pathway.

Several simplifying assumptions are used to model the hybrid system:

i. Spatial gradients in the biofilm were neglected (zero spatial dimension);
ii. Transport limitation is considered the dominant process in the biofilm (maximum AMX process rate, *ρ_AMX,max_*, and AMX biomass concentration, *X*_AMX_, are constant);
iii. No explicit oxygen inhibition of AMX (reflected in the maximum volumetric AMX activity constant, *r*_AMX,max_ g_N_m^-3^d^-1^);
iv. Perfect biomass segregation between floc (AOB, NOB) and biofilm (AMX) functional guilds;
v. Decay is considered negligible and ignored;
vi. Quasi steady-state assumptions describing oxygen concentration and non-aeration SBR phases.

These allow for a conventional dynamic model to be developed for simulating the PN/A process under different operating strategies with respect to three control parameters; dissolved oxygen concentration (S_O2_ g_O2_ m^-3^), fraction of solids removed (*f*_was_), and maximum Anammox activity. Item vi describe critical simplifying assumptions by which oxygen concentration in the bulk liquid phase is assumed constant (i.e. ideal control at fixed oxygen set-point) and the decanting, mixing, and refilling steps of the SBR process occur instantaneously. In other words, these actions are considered to occur at a much faster rate than the dynamics of the system state variables.

Whilst the simple model may not provide quantitative predictions that align with experimental data, qualitative correspondence was demonstrated by the authors. With this in mind, here we show that numerical analysis of the model as an impulsive system of ODEs gives a more complete insight into the behaviour of the process. Given that the results of the analysis are still constrained by the simplicity of the model they, nevertheless, provide a more detailed picture of the SBR operating space with respect to the operating parameters, and the methodology also allows for analysis to be extended to any of the model parameters. This provides opportunities to investigate the impact of guild interactions on process performance in a more consistent and systematic manner.

## 2 The impulsive model

We reformulate the model first presented by Laureni *et al.* [16] explicitly as an impulsive system. Given the standard notation for impulsive differential equations (see Appendix A.1) then, for a function *y*(*t*) and time *τ*,

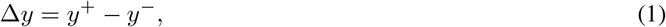

where

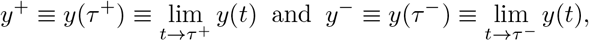

we define a system of equations (adapted from [16], with notations given in Appendix A.2) describing the aeration and refilling steps of the sequencing batch operation:

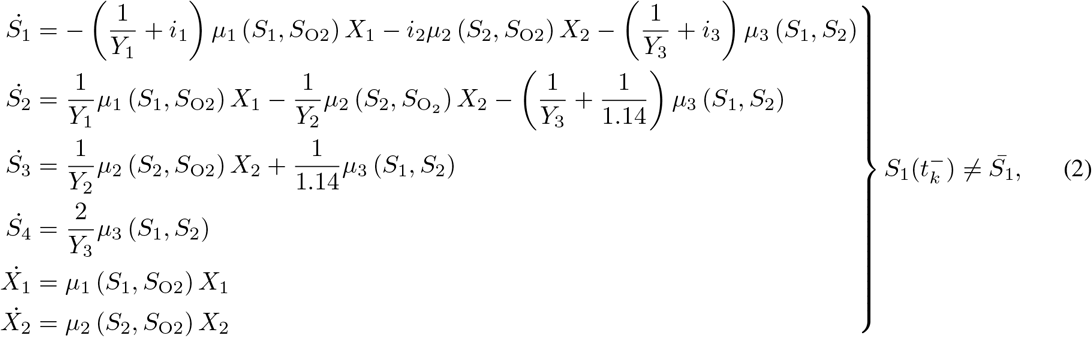

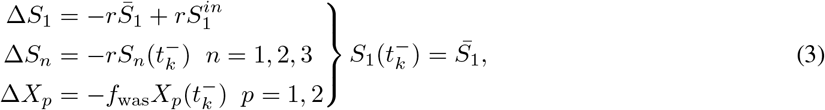

where *μ_i_* (*i* = 1,2,3) are the monotonically increasing Monod growth functions, *r* is the refill fraction, *f*_was_ is the fraction of solids wasted at the end of each cycle, 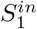 is the ammonium added to the reactor at the start of each cycle, and 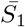 is the threshold ammonium concentration that triggers the end of the aeration phase and the subsequent refilling step. The maximum Anammox process rate (*ρ*_3_ ≡ *μ*_*m*,3_*X*_3_), which is assumed constant, is given by

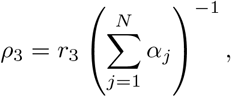

where *r*_3_ is the maximum volumetric Anammox activity and *α_j_, j* ∈ [*S*_1_, *S*_2_], are the stoichiometric coefficients associated with the ammonium and nitrite consumption by the Anammox process. The variable growth terms describing multiple substrate Monod type functions are given by:

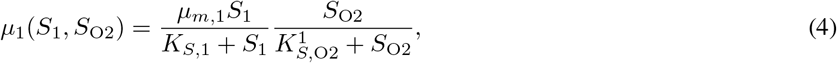

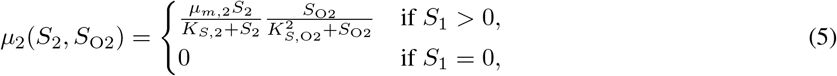

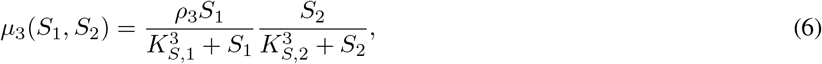

where *ρ*_3_ ≡ *μ*_3_*X*_3_ is the maximum Anammox process rate assuming constant Anammox biofilm density. Note that NOB growth rate (Eq. 5) is zero if ammonium concentration is also zero, a conditional check to avoid a negative reaction rate in S1 [25, 10].

Denoting *t_k_* as the cycle time for which the ammonium concentration reaches the threshold value, 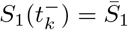, then from equations (1) and (3), the component concentrations at the start of each SBR cycle are given by:

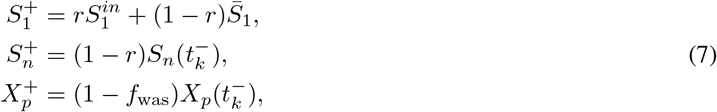

where 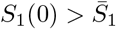.

### 2.1 Performance metrics

The objective of the study by Laureni *et al*. [16], to understand the qualitative behaviour of the competing nitrifiers under differential operating conditions, is underpinned by identification of the ideal operating scenario for complete NOB washout. These are predicated on the model assumptions above, particularly Items ii - iv. Thus, we can describe how a transcritical bifurcation (denoted hereafter as a branch-point) separates a two parameter phase space into ideal and non-ideal operating regions. A transcritical bifurcation is characterised as a fixed-point, or equilibrium, whose stability interchanges with the stability of another fixed-point as the bifurcation parameter is varied. In our case, we have the interior equilibrium, 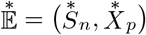, interchanging with the NOB washout equilibrium, 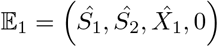, and a branch-point curve can be drawn as the bisection of these two regions as a two parameter bifurcation diagram.

To assess the performance of the reactor, nitrogen removal rate and efficiency are calculated over the SBR cycle. This indicates the degree of residual NO_x species in the reactor after the ammonium threshold concentration (2 mg_NH4-N_L^-1^) has been met. The Hydraulic Retention Time (HRT) is equivalent to the cycle time (*T*) under simplifying assumptions (AOB biomass concentration and growth are approximately constant over a single cycle), and is dependent on flocs wastage by *f*_was_ ≈ *μ*_1_*T* [16].

Nitrogen removal efficiency (%) is calculated by:

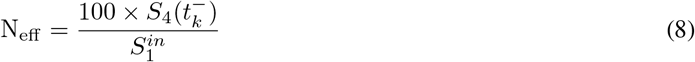

Hydraulic Retention Time (hours) per cycle is calculated by:

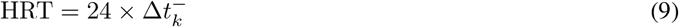

given 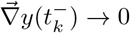, where *y* denotes the system of equations (2). This condition is an assumption that the impulsive system reaches some fixed periodic orbit for *t_k_* large enough, but in practice we wish to approximate convergence to avoid long simulation times. Thus, we consider a fixed periodic orbit is reached when the maximum biomass concentration gradient, 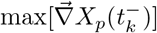, falls below a tolerance equivalent to 1% of the norm of its absolute value, 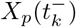.

The magnitude of the two performance indicators can highlight suitable operating conditions for good process performance, for example high nitrogen removal efficiency at low HRT. By performing the bifurcation analysis of the impulsive system, process performance can be assessed directly in relation to steady-state behaviour, providing a richness of information absent through simulation alone.

### 2.2 Analysis

Bifurcation analysis is an important method for gaining critical insights into systems of continuous or discrete ODEs, describing their behaviour in relation to key control parameters. The impulsive system given by Equations (2) and (3) consists of both continuous (the reaction phase) and discrete (the decant-refill step) time components that require tools that can perform bifurcation analysis of iterated maps. Importantly, we require a software that is able to perform continuation of the fixed points of an iterated map in two parameters. The continuation software `CL_MatContM` [5] is utilised here as it provides the necessary tools for identifying codimension-1 bifurcations and allows for the continuation in two dimensions of the impulsive map. Although the `MatContM` code is written for use in the proprietary software `MATLAB` (version 9.6.0, The MathWorks, Inc., Natick, Massachusetts, United States), the command line (CL) version is readily modified to work with impulsive maps, which is not possible via the Graphical User Interface.

We start by finding the values of the six model variables at the interior equilibrium ([*S_n_, X_p_*] > 0). To do this, we simulate the system using the open-source numerical solver software `XPP` [3] with initial conditions [*S*_1_ = 20, *S*_2_ = 0,*X*_1_ = 5.6,*X*_2_ = 12.5] (gm^-3^) and operating parameters [*S*_O2_ = 1.5 g_O2_m^-3^, r_AMX,max_ = 86 g_N_m^-3^, f_was_ = 0.5 %] (see `DM5_impulsive_paper.ode` file in Supplementary Software). Note that *S*_3_ and *S*_4_ are not included in the simulation as they decouple from the model and are not required for identification of equilibria. We check, with *t* large enough, that the system converges to the interior equilibrium 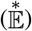 by inspecting the values at a section of the Poincaré map intersecting the periodic orbits. That is, as the system settles to a constant period, we choose an admissible value in one variable (i.e. *S*_1_ = 4.0) and note the magnitude ofthe other variables as the periodic orbit for *S*_1_ passes through this value. When these variables at the section converge to within a given tolerance (*ϵ* < 1%), we note their values at the subsequent impulse time 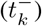 and stop simulating. These steps are summarised in Procedure 1.

#### Procedure 1 Find fixed periodic orbit

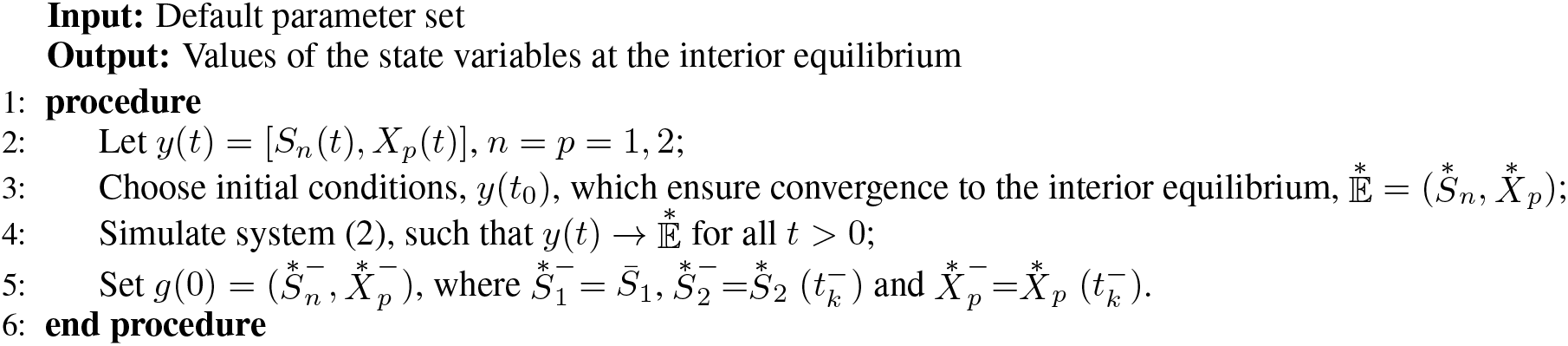

We can now use the fixed-points as our initial data, *g*(0), for bifurcation analysis using `CL_MatContM` [6]. Selecting *P_a_* ∈ [*S*_O2_, r_AMX,max_, *f*_was_] to be active bifurcation parameter, we first ensure the solution is at a fixed-point by performing numerical continuation in *P_a_* to assure convergence. In the instances where multiple bifurcation points are detected, we select only the branch-point separating the interior and the NOB washout equilibria. We can now follow the branch-point in two-parameter space to produce a curve that separates these two regions of equilibria. Although a transcritical bifurcation (branch-point) is more degenerate than a limit point bifurcation (also called a fold bifurcation or saddle-node bifurcation), in our case, the conditions for their emergence are taken as equivalent, where the only qualitative difference is that in the former, there are two equilibria either side of the critical bifurcation point (with corresponding change of stability along the branches) and, in the former, two equilibria fall on one side of this value but none on the other. This allows us to use the limit point map for continuation of the branch-point curves, as all conditions for identification of these points are met under the equivalency.

We use the *limitpointmap* function from `CL_MatContM` to continue the branch-point forward and backward in two-parameter space, [*P_a_*, *P_b_*]. Procedure 2 provides the steps for producing these curves and the `CL_MatContM` code, which can be executed in the `MATLAB` environment, is provided in the Supplementary Software (`branch_curves.m`).

#### Procedure 2 Follow Branch-Point Curve in two-parameters

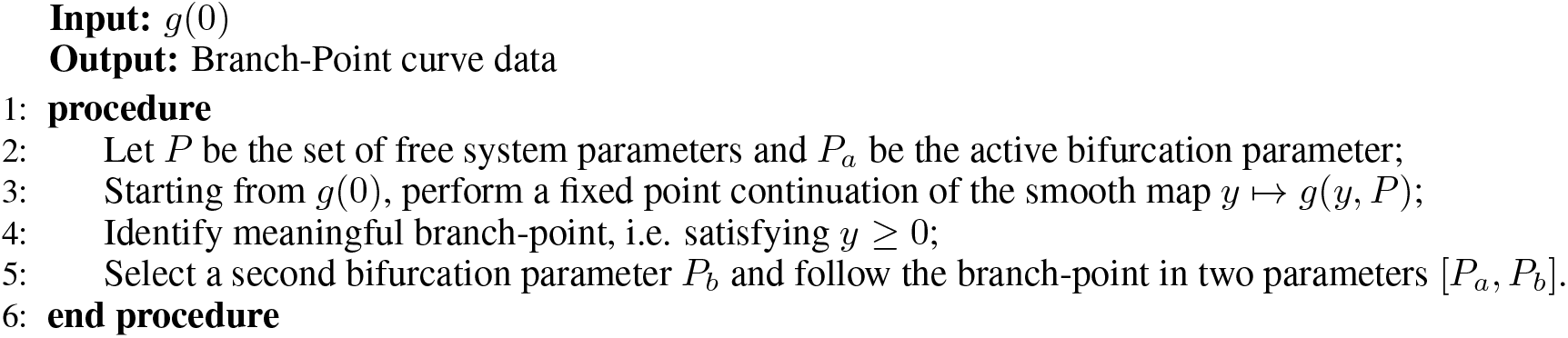

## 3 Results and discussion

We performed bifurcation analysis of the impulsive system defined by the model described by Laureni *et al*. [16]. In a first step, we simulated the system with initial conditions and parameters (see Section 2.2) that guaranteed convergence to the interior equilibrium, such that AOB and NOB populations are non-zero. From this fixed-point we then performed a one-parameter bifurcation using dissolved oxygen or the AMX activity constant, which revealed two branch-point bifurcations in the admissible region (i.e., [*S_n_*, *X_p_*] ≥ 0). These branch-points relate to transitions between the three possible system equilibria:

- 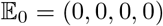 - AOB and NOB washout;
- 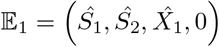 - NOB washout;
- 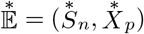 - AOB and NOB coexist.

Given that the principle operating objective of PN/A systems, particularly under mainstream conditions, is the suppression of NOBs [15, 16], then we focus on the branch-point occurring when parameter variations result in the transition from 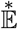 to 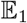. Hence, we developed bifurcation diagrams by following this branch-point in two parameters superimposed onto contour plots showing the Nitrogen removal efficiency, see Figs 1 (a), (c), (e), and HRT, see Figs 1(b), (d), (f). The branch-point was continued both in forward and backward directions to obtain the smooth branch-point curves indicated. We confine the extent of the parameter variation to approximately those given in the original empirical work [16]. The figures are also marked with the points representing the parameter values investigated by Laureni *et al*. [16], with the third (fixed) operating parameter set to its default value when not used as a bifurcation parameter. The single black marker indicates default operation where AOB and NOB coexist, whilst the white markers are the represent a change in one control parameter that facilitates NOB washout.

**Figure 1:**
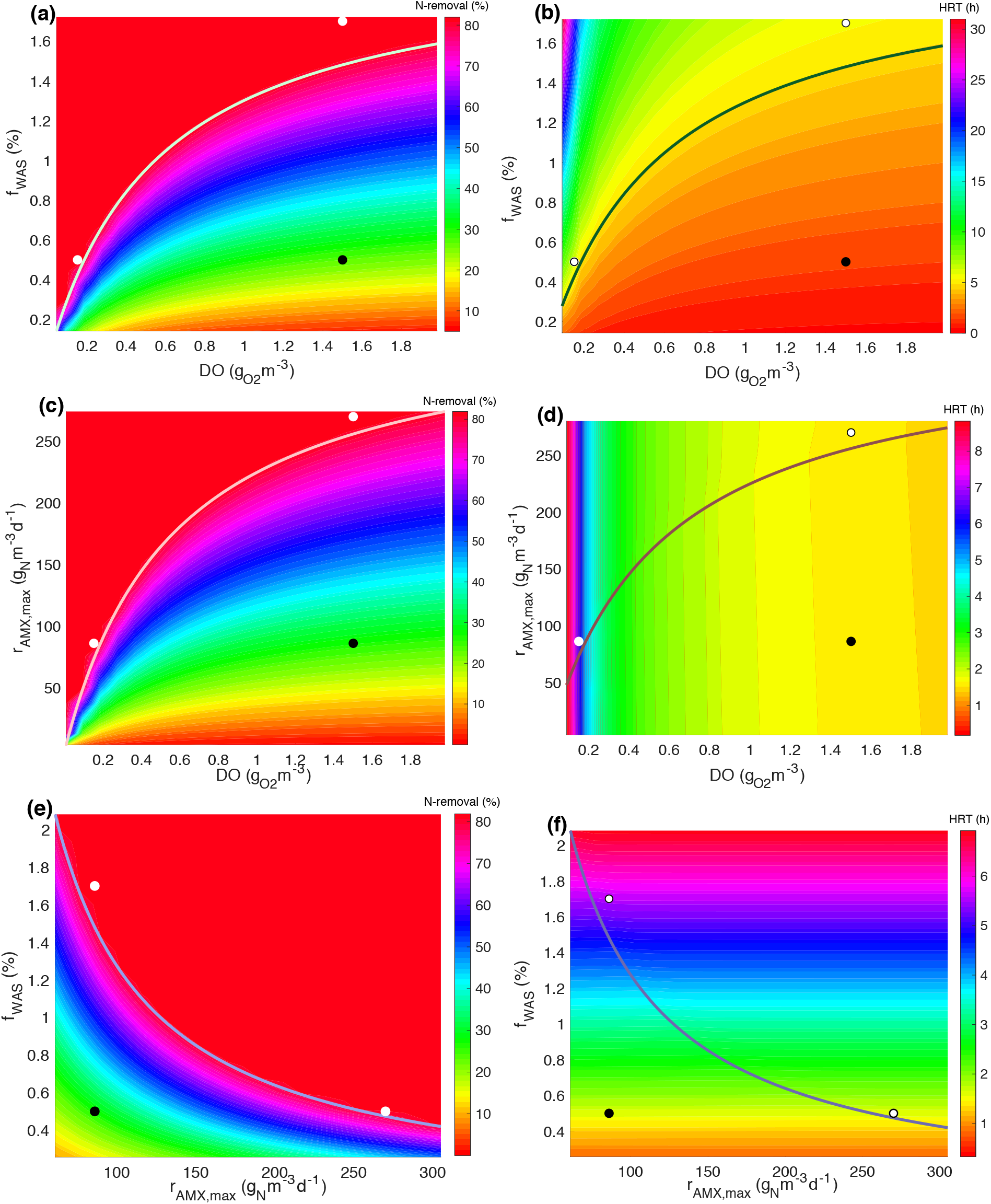
Diagrams showing two-parameter bifurcations describing the branch-point curves separating the region where two species (AOB and NOB) exist and where only the AOB exists (NOB is washed out). The black points are the default operating values given by Laureni et al. [16] and lie in the former region, and the white dots signify single parameter changes moving the process to the NOB washout equilibria. Contour plots are added to the figures to indicate magnitudes of two performance indicators within the parameter space; Left - Nitrogen removal efficiency (%), Right - Hydraulic Retention Time (h).

The results show that the branch-point curves bisect a region of nearly constant nitrogen removal efficiency 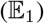 and a region with graduated removal dependent on operating parameters. This corresponds with observations from other studies indicating that a decrease in oxygen concentration, an increase in either floc wastage or Anammox activity will result in conditions in which NOB diminish. However, here was can demonstrate the theoretical three parameter operating domain in which to achieve this objective.

The figures showing these bifurcation curves against HRT indicate a decoupling between relative AOB-NOB activity and cycle time. Instead, the HRT can be viewed as a potential control objective in relation to the operating parameters, and identification of operational ‘sweet-spots’ may be assessed accordingly. For example, we can observe that under default conditions, we can obtain maximum nitrogen removal efficiency by lowering DO to 0.18 g_O2_m^-3^ and this results in an HRT of approximately 6 hours, a value shown to be optimal in similar empirical SBR studies, albeit at elevated temperatures [31, 17]. However, a further reduction in oxygen increases the HRT considerably. Similarly, increasing the fraction of floc wastage to 1.7 % results in a similar HRT, but in tandem with DO reduction, increases HRT to approximately 20 hours. The effect of Anammox activity on HRT is minor, as was observed in the original study (see Scenario 4 in [16]).

Whilst management of Anammox biofilm activity is theoretically possible via enrichment or modified attachment rates [13], oxygen control and floc removal are the more reliable methods to achieve selective NOB washout. Given a low Anammox activity equivalent to a young biofilm, for example, and relatively high SRT (e.g. low *f*_was_), operating at low oxygen concentrations appears sufficient to guarantee maximum nitrogen removal. However, it can be seen in the figure that operation is close to the theoretical limit for NOB washout and, given the simplifying assumptions of the model, operating at lower SRT may account for any uncertainty in Anammox activity rates.

In practice it is implausible that complete NOB removal can be achieved. Ultimately, the mode of operation will be dependent on process objectives. The use of sequencing batch operation ensures a fixed effluent ammonium concentration and secondary targets such as nitrogen removal efficiency and cycle time can be guided by simulation and bifurcation analysis, as demonstrated here. It can be noted that good nitrogen removal efficiencies (70-80 %) occur when NOB are present in the reactor. Many empirical studies focus on trying to minimise NOB populations and their regrowth, which is not trivial. Whilst the model presented here has a number of simplifying assumptions, the analysis could be used to investigate process behaviour when NOB are not entirely removed from the system, e.g. allowing for some denitrification by inclusion of heterotrophs in the model.

## 4 Conclusions

Mathematical analysis of impulsive systems is non-trivial in many cases, but for engineers numerical studies provide sufficient information. Early attempts at characterising biological nitrogen removal in a SBR using bifurcation analysis were restricted to continuous ODEs describing growth of a single organism [26]. Analytical studies of nitrification by the coupling of AOB and NOB (e.g. the SHARON process) have facilitated general mathematical description of these systems indicating the possibility for multiplicity of equilibria and their stability [23, 24, 25]. However, these systems have assumed continuous operation, meaning the models are tractable to analysis using standard mathematical tools. Similarly, a rigorous mathematical analysis of a two species partial nitritation reactor using different dilution rates to mimic biomass retention, has recently shown the ability for mathematics to further engineering application, in this case through dissolved oxygen control [10].

Here, we have performed numerical analysis of a three species partial nitritation/Anammox process operated as a SBR. Building on the recent model developed by Laureni *et al*. [16], we implemented a novel method for following a branchpoint equilibrium describing the boundary between ideal and non-ideal conditions for three identified control parameters. Coupled with data describing key performance indicators for assigned control parameter pairs, the bifurcation analysis provides both a greater level of knowledge regarding operating regimes, but also a method to investigate the influence of system configuration and operation on reactor behaviour, in general.

Indeed, given the overtly simplified model used in this study, the opportunities to investigate the influence of factors not considered here becomes a primary objective for further work. For example, the effect of including heterotrophs or polyphosphate-accumulating organisms, which compete with the nitrifiers for resources, may reveal unexpected behaviour given the increased ‘complexity’ of the modelled system.

Ultimately, this study has reinforced the idea that mathematical analysis has an important role to play in the understanding and control of engineered biological systems. Whilst the level of detail and the simplifying assumptions of the underlying model may reduce the ability to predict behaviour with quantitative accuracy, qualitative insights are vital to understanding open questions and future directions for biological treatment processes in a time when resource and energy efficiency is such a prominent focus.

## Acknowledgments

M.J.W. acknowledges the support from the European Union’s Horizon 2020 research and innovation programme under the Marie Skłodowska-Curie Grant Agreement No. 702408 (DRAMATIC).

## Appendix A Notations and parameters

### A.1 Impulsive system notations

We describe the standard notation of impulsive systems used here in lay terms. Assuming the standard definition of a differential equation:

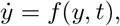

and the moments of the impulsive effect for its solution, *y*(*t*), occur when *t* = *τ_k_*, where *k* ∈ *N* is the index in a sequence of impulses. Given the quasi steady-state assumption from Item vi, we can define an impulsive map, *I*(*t, y*), which is a function that maps the system solution before the impulse, 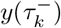, to the solution after the impulse, 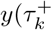, such that:

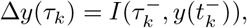

and 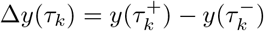 (see Eq. 1). For consistency, we represent the impulsive time by *t_k_* ≡ *τ_k_*.

### A.2 Model notations

We use here a parsimonious notation adapted from that given by Laureni *et al.* [16]. State variables *S_n_* and *X_p_* use subscript integers rather than subscript characters representing metabolites and species, except in the case of oxygen where we maintain the character representation as it is assumed constant here. Table A.2 provides the original and new notations for the state variables for ease of comparison, whilst Table A.2 gives the default model parameters and their values.

**Table 1:** Original and new notations for model state variables

| Original notation [16] | New notation | Description |
| --- | --- | --- |
| $S_{\text{NH}_4}$ | $S_1$ | Ammonium concentration ($\text{g}_\text{N}\text{m}^{-3}$) |
| $S_{\text{NO}_2}$ | $S_2$ | Nitrite concentration ($\text{g}_\text{N}\text{m}^{-3}$) |
| $S_{\text{NO}_3}$ | $S_3$ | Nitrate concentration ($\text{g}_\text{N}\text{m}^{-3}$) |
| $S_{\text{N}_2}$ | $S_4$ | Dinitrogen gas concentration ($\text{g}_\text{N}\text{m}^{-3}$) |
| $S_{\text{O}_2}$ | $S_{\text{O}_2}$ | Oxygen concentration ($\text{g}_{\text{O}_2}\text{m}^{-3}$) |
| $X_{\text{AOB}}$ | $X_1$ | AOB biomass concentration ($\text{g}_{\text{COD}}\text{m}^{-3}$) |
| $X_{\text{NOB}}$ | $X_2$ | NOB biomass concentration ($\text{g}_{\text{COD}}\text{m}^{-3}$) |
| $X_{\text{AMX}}$ | $X_3$ | Anammox biomass concentration ($\text{g}_{\text{COD}}\text{m}^{-3}$) |

**Table 2:** Kinetic, stoichiometric and operating parameters used in the impulsive model

| Parameter | Description | Value | Unit |
| --- | --- | --- | --- |
| $\mu_{m,1}$ | Maximum specific growth rate (AOB) | 0.30 | $\text{d}^{-1}$ |
| $K_{S,1}$ | Ammonium half-saturation constant (AOB) | 2.4 | $\text{g}_{\text{NH}_4\text{-N}}\text{m}^{-3}$ |
| $K_{S,\text{O}_2}^1$ | Oxygen half-saturation constant (AOB) | 0.6 | $\text{g}_{\text{COD}}\text{m}^{-3}$ |
| $Y_1$ | Growth Yield (AOB) | 0.18 | $\text{g}_{\text{COD}}\text{g}_\text{N}^{-1}$ |
| $i_1$ | Nitrogen required for cell synthesis (AOB) | 0.083 | $\text{g}_\text{N}\text{g}_{\text{COD}}^{-1}$ |
| $\mu_{m,2}$ | Maximum specific growth rate (NOB) | 0.34 | $\text{d}^{-1}$ |
| $K_{S,2}$ | Nitrite half-saturation constant (NOB) | 0.5 | $\text{g}_{\text{NO}_2\text{-N}}\text{m}^{-3}$ |
| $K_{S,\text{O}_2}^2$ | Oxygen half-saturation constant (NOB) | 0.4 | $\text{g}_{\text{COD}}\text{m}^{-3}$ |
| $Y_2$ | Growth Yield (NOB) | 0.08 | $\text{g}_{\text{COD}}\text{g}_\text{N}^{-1}$ |
| $i_2$ | Nitrogen required for cell synthesis (NOB) | 0.083 | $\text{g}_\text{N}\text{g}_{\text{COD}}^{-1}$ |
| $K_{S,1}^3$ | Ammonium half-saturation constant (AMX) | 0.03 | $\text{g}_{\text{NH}_4\text{-N}}\text{m}^{-3}$ |
| $K_{S,2}^3$ | Nitrite half-saturation constant (AMX) | 0.005 | $\text{g}_{\text{NO}_2\text{-N}}\text{m}^{-3}$ |
| $Y_3$ | Growth Yield (AMX) | 0.17 | $\text{g}_{\text{COD}}\text{g}_\text{N}^{-1}$ |
| $i_3$ | Nitrogen required for cell synthesis (AMX) | 0.058 | $\text{g}_\text{N}\text{g}_{\text{COD}}^{-1}$ |
| $S_1^{\text{in}}$ | Influent ammonium concentration | 20 | $\text{g}_{\text{NH}_4\text{-N}}\text{m}^{-3}$ |
| $\bar{S}_1$ | Threshold ammonium concentration | 2 | $\text{g}_{\text{NH}_4\text{-N}}\text{m}^{-3}$ |
| $\rho_3$ | Anammox process rate | varied | $\text{g}_{\text{COD}}\text{m}^{-3}\text{d}^{-1}$ |
| $r$ | Volumetric exchange fraction per SBR cycle | 0.5 | - |
| $f_{\text{was}}$ | Floc removal fraction per SBR cycle | varied | - |

